# Pan-cancer analysis identifies nine conserved miRNA regulators of tumor cytolytic activity and clinically actionable immune targets

**DOI:** 10.64898/2026.08.30.748071

**Authors:** Nazanin Bagherlou, Shahram Aliyari, Zahra Salehi, Mobin Pirouzkhah, Cleo-Aron Weis

## Abstract

**Background:** Cytolytic activity (CYT), a widely used transcriptomic surrogate of anti-tumor immune cytotoxicity derived from GZMA (granzyme A) and PRF1 (Perforin 1) expression, is associated with clinical outcomes across cancers. MicroRNAs (miRNAs) are key post-transcriptional regulators of tumor immunity, yet their pan-cancer roles in modulating cytolytic activity remain incompletely understood.

**Objective:** This study aimed to identify conserved miRNA regulators of tumor cytolytic activity and their downstream gene-mediated networks across diverse cancer types, while evaluating their clinical and therapeutic relevance.

**Methods:** Matched miRNA and mRNA expression profiles from 9,288 primary tumors across 31 TCGA cancer types were analyzed. A multi-stage framework was applied: per-cancer Spearman correlations (|ρ| ≥ 0.30, FDR < 0.05) identified recurrent CYT-associated miRNAs (≥3 cancer types); these were integrated with TargetScan-predicted targets and subjected to pan-cancer and cross-cancer triple filtering (miRNA–gene and gene–CYT associations). All associations underwent tumor purity adjustment using Consensus Purity Estimate (CPE), with LUMP (Leukocytes Unmethylation for Purity) as sensitivity analysis. Candidates were further prioritized by random forest modeling with bootstrap stability, cancer-type-adjusted Cox regression, mediation analysis, immune cell deconvolution, k-means molecular subtyping, pathway enrichment, and DGIdb-based drug-target prioritization.

**Results:** The analysis converged on 38 high-confidence miRNA–gene–CYT regulatory triplets involving 9 conserved miRNAs and 31 target genes after stringent purity adjustment and multi-layer validation. All nine miRNAs exhibited complete bootstrap stability. Mediation analysis confirmed significant gene-level mediation in 37 of 38 triplets (FDR < 0.01), with mediated proportions up to 94%. The final miRNA signature defined two distinct pan-cancer immune subtypes (immune-hot vs. immune-cold) with significantly different cytolytic activity and overall survival (OS) (HR = 0.754, FDR = 1.12 × 10⁻⁴). The network was enriched for T-cell activation and lymphocyte differentiation pathways and highlighted multiple druggable targets, including CTLA4 and CD274 (PD-L1), nominating 124 candidate compounds.

**Conclusions:** In conclusion, this tumor purity-adjusted pan-cancer study defines a compact, reproducible, and clinically relevant miRNA network that regulates cytolytic activity across diverse malignancies. By linking miRNA biology to immune subtyping and actionable therapeutic targets, the present work provides a valuable foundation for advancing precision immuno-oncology.

## Introduction

Cytolytic activity (CYT), commonly quantified as the arithmetic mean of log₂-transformed GZMA (granzyme A) and PRF1 (Perforin 1) expression, is a widely used transcriptomic surrogate of anti-tumor immune-mediated killing and reflects the activity of cytotoxic T lymphocytes and natural killer cells within the tumor microenvironment (TME) (1, 2). Across many cancer types, elevated CYT has been associated with improved patient survival, enhanced anti-tumor immunity, and favorable responses to immunotherapy (3–5). However, the biological interpretation of CYT is context-dependent. In certain tumors, including gliomas, high CYT may coexist with immune exhaustion, chronic inflammation, and immunosuppressive microenvironmental features, resulting in paradoxically unfavorable clinical outcomes (6, 7). These observations suggest that understanding the molecular regulators of CYT is critical to deciphering the mechanisms of effective versus dysfunctional anti-tumor immunity.

MicroRNAs (miRNAs) are small non-coding RNAs of approximately 22 nucleotides that regulate gene expression primarily through interactions with target mRNAs, resulting in translational repression or transcript degradation (8). Within the immune system, miRNAs regulate hematopoietic development, immune cell differentiation, activation, and effector functions, thereby shaping immune responses during infection, aging, and malignant transformation (9). Aberrant miRNA expression is a hallmark of many cancers and contributes to tumor progression, immune escape, and TME remodeling (8). Although miRNAs are classically regarded as negative regulators of gene expression, accumulating evidence indicates that they can also exert indirect or context-dependent positive regulatory effects through complex cellular networks (10, 11).

Several miRNAs have already been implicated in pathways relevant to cytolytic immunity. Among these, hsa-miR-155-5p functions as a pleiotropic regulator of immune activation and lymphocyte responses (12–14), whereas members of the miR-29 and miR-200 families have been linked to tumor-suppressive, stromal, and immune-modulatory programs (15, 16). Nevertheless, the broader landscape through which miRNAs influence CYT across diverse cancer types remains poorly understood.

Most computational studies investigating miRNA–CYT relationships rely on pairwise correlation analyses that evaluate associations between miRNAs and immune phenotypes independently (17). Such approaches are limited because they fail to account for the intermediate gene regulatory mechanisms through which miRNAs are expected to influence immune function. Moreover, the assumption that biologically meaningful miRNA effects must always involve inverse miRNA–gene relationships overlook accumulating evidence for indirect and positive regulatory associations observed in cancer systems (10, 11, 18). As a result, many potentially relevant miRNA-mediated immune regulatory networks may remain undetected. To date, no pan-cancer study has systematically integrated miRNA–CYT associations with target-gene relationships, mediation analysis, tumor purity correction, cross-cancer recurrence assessment, machine-learning prioritization, and immune subtype discovery within a unified analytical framework.

In this study, we developed a multi-layered pan-cancer framework to identify recurrent miRNA-associated regulators of cytolytic anti-tumor immunity across 32 cancer types from The Cancer Genome Atlas (TCGA). We hypothesized that a subset of miRNAs exerts consistent associations with CYT through gene-mediated regulatory pathways that are conserved across multiple tumor types and possess prognostic relevance. To test this hypothesis, we implemented a triangular regulatory model requiring simultaneous evidence of significant miRNA–CYT, miRNA–gene, and gene–CYT associations. Both negative and positive miRNA–gene co-variation patterns were considered, and all associations were re-evaluated after adjustment for tumor purity. Candidate regulators were subsequently prioritized using random forest–based importance and stability metrics, evaluated for prognostic significance using cancer-type–adjusted survival analyses, examined through formal mediation testing, and further characterized by immune microenvironment profiling and unsupervised pan-cancer subtype discovery. Through this strategy, we aimed to generate a comprehensive map of miRNA-associated regulatory programs linked to cytolytic immunity and to identify candidate biomarkers and therapeutic targets for future immuno-oncology research (Fig. 1).

**Figure 1.**
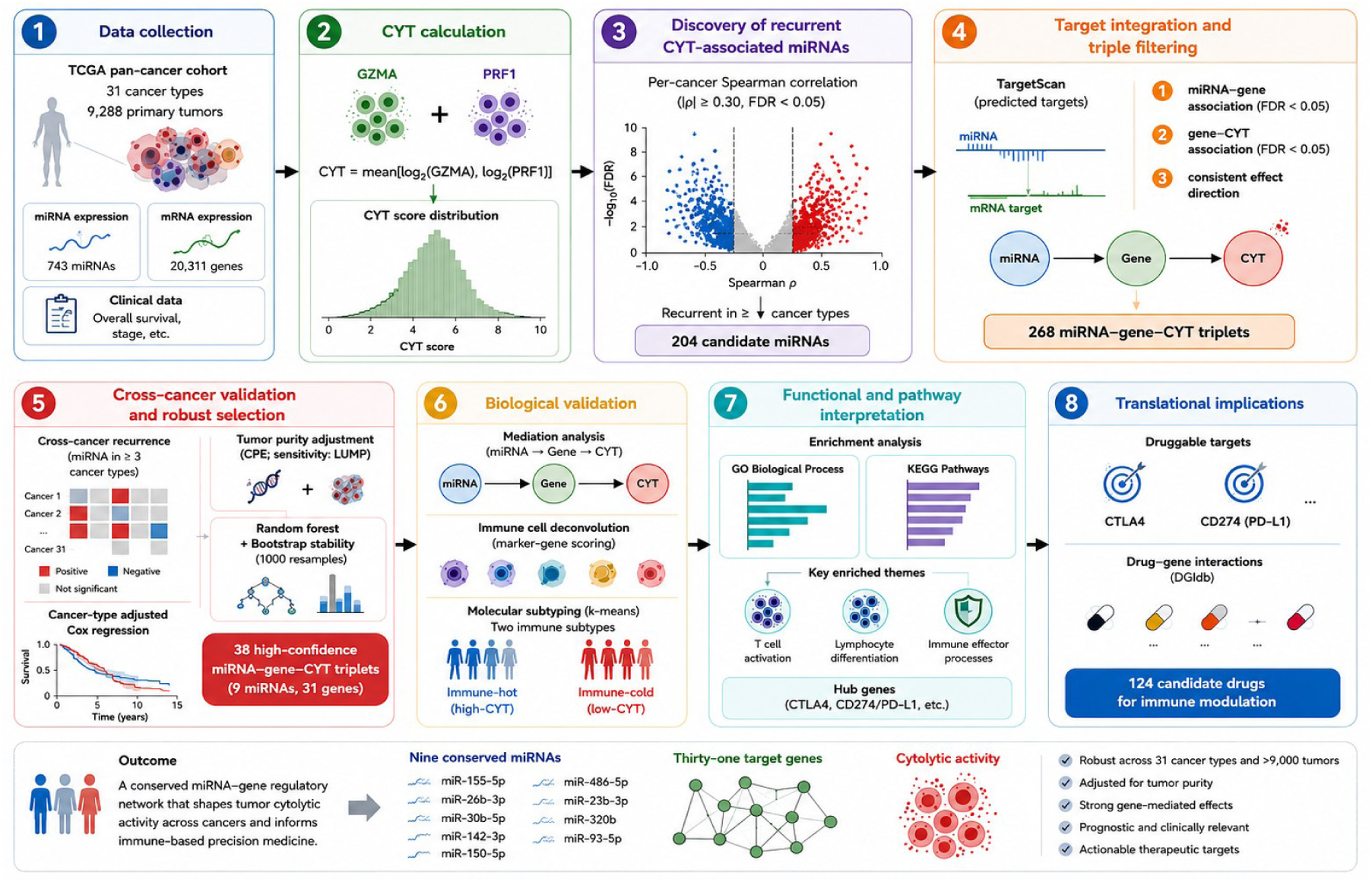
Overview of the analytical workflow for identifying pan-cancer miRNA–gene regulatory networks associated with CYT. Primary tumour samples from 31 TCGA cancer types (n = 9,288) were integrated with miRNA, mRNA, and clinical data. CYT was calculated from the log₂-transformed expression of *GZMA* and *PRF1*. Candidate miRNAs were identified by cancer-specific Spearman correlation analyses with CYT, followed by TargetScan-based miRNA–gene integration and a three-step filtering strategy requiring significant miRNA–gene association, significant gene–CYT association, and concordant regulatory direction. Recurrent associations detected in at least three cancer types were subsequently refined by tumour-purity adjustment (CPE, with LUMP sensitivity analysis), machine-learning prioritization using random forest with bootstrap stability assessment, and cancer-type-adjusted survival analysis. High-confidence miRNA–gene–CYT triplets were further evaluated by mediation analysis, immune-cell association analysis, molecular subtype discovery, pathway enrichment, and drug–target prioritization, resulting in a robust set of recurrent immune-associated regulatory networks with potential translational relevance.

## Materials and Methods

### Data Sources

mRNA expression data were obtained from the TCGA pan-cancer empirical Bayes-adjusted (EB-adjusted) RNA-SeqV2 dataset available through the UCSC Xena Pan-Cancer Hub, which provides log₂-normalized, batch-corrected expression values (19). miRNA expression data were obtained from the corresponding TCGA pan-cancer EB-adjusted miRNA expression dataset available through the same platform. Clinical data, including overall survival (OS) time and event status, were obtained from the Pan-Cancer Clinical Data Resource (20). Tumour purity estimates were retrieved using the TCGAbiolinks R package (21), including the Consensus Purity Estimate (CPE) and Leukocytes Unmethylation for Purity (LUMP) metrics. Predicted miRNA–target gene interactions were obtained from TargetScan (22) using context++ scores and were restricted to human targets. Experimentally validated miRNA–target interactions were obtained from mirtarbase (23), using both the complete set of documented interactions and a subset restricted to strong experimental evidence (reporter assays and Western blotting). Drug– gene interaction data were obtained from the Drug–Gene Interaction Database (DGIdb) (24).

### Cohort Assembly and Sample Harmonization

TCGA barcodes were standardized to a 15-character format and restricted to primary tumour samples (sample type code 01). Samples were matched across four data layers: miRNA expression, mRNA expression, CYT score, and survival data, using exact sample identifiers. The matched analytical cohort comprised 9,288 primary tumour samples spanning 31 cancer types. Glioblastoma multiforme (TCGA-GBM) was excluded because matched mature miRNA data were unavailable in the harmonized dataset. When cancer type annotations were unavailable in the Pan-Cancer Clinical Data Resource, cancer type was inferred from the TCGA project code contained within the sample barcode. Cancer types were included in per-cancer analyses only if they contained at least 30 matched samples. Following quality control, the final matched expression matrices comprised 743 miRNA features and 20,311 mRNA features. Prior to association analyses, both expression matrices underwent uniform preprocessing as described below.

### Normalization and Quality Control

Both the miRNA and mRNA expression matrices were verified to be log₂-transformed prior to analysis; therefore, no additional logarithmic transformation was applied. Expression values were winsorized for each feature at the 0.1th and 99.9th percentiles to limit the influence of extreme values, and features exhibiting zero variance across all samples were excluded. For correlation analyses, each feature was required to have at least three finite observations and non-zero variance across samples.

### Cytolytic Activity Score

CYT score was derived from the expression of GZMA and PRF1, two established markers of cytotoxic lymphocyte activity originally proposed by Rooney et al.

For sample *i*, CYT was calculated as:

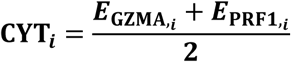

where 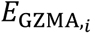 and 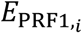 denote the log₂-normalised expression values of GZMA and PRF1, respectively.

This formulation represents the arithmetic mean of two log₂-normalised expression values and is conceptually related to the original CYT metric described by Rooney et al., which was calculated as the geometric mean of GZMA and PRF1 expression on the original non-logarithmic scale. Consequently, absolute CYT values are not directly comparable between studies; however, the present formulation preserves relative sample ranking and is appropriate for the rank-based and association analyses performed in this study. CYT was modelled as a continuous outcome variable in all subsequent correlation, tumour purity-adjusted, machine-learning, mediation, subtype, and survival analyses.

### Identification of Candidate miRNA–Gene–CYT Regulatory Triplets

Within each eligible cancer type (n ≥ 30 matched samples), Spearman rank correlation (25) was computed between each of the 743 miRNA features and CYT. Multiple testing was controlled using the Benjamini–Hochberg (BH) procedure to control the false discovery rate (FDR) (26), applied independently within each cancer type. A miRNA was classified as significantly associated with CYT within a given cancer type if FDR < 0.05 and |ρ| ≥ 0.30. Candidate miRNAs were defined as those meeting both criteria independently in at least three cancer types. These candidates formed the basis for subsequent integration with predicted miRNA–target gene interactions.

TargetScan-predicted miRNA–target gene interactions were integrated with the matched expression matrices for downstream analyses. For each candidate miRNA and each predicted target gene present in the matched mRNA expression matrix, two pan-cancer Spearman correlations were computed across all 9,288 samples: (i) miRNA expression versus target gene expression and (ii) target gene expression versus CYT. BH correction was applied jointly across all miRNA–gene correlation tests and separately across all gene– CYT correlation tests. A miRNA–gene–CYT triplet was retained if both correlations satisfied the predefined effect size and significance criteria (|ρ| ≥ 0.30 and FDR < 0.05). Retained triplets were classified into directional branches according to the signs of the corresponding Spearman correlation coefficients. The negative branch comprised triplets with ρ (miRNA, gene) < 0, whereas the positive branch comprised triplets with ρ (miRNA, gene) > 0. Triplets passing the pan-cancer filtering criteria were subsequently evaluated for consistency across individual cancer types.

Each miRNA–gene pair passing the pan-cancer triple filter was re-evaluated independently within each eligible cancer type using the predefined correlation and significance thresholds for both the miRNA–gene and gene–CYT associations. Pairs were retained for downstream analyses only if they were independently detected in at least three cancer types.

For visual representation of recurrent miRNA–gene interactions, an edge score was assigned to each retained pair based on the strength of both pan-cancer association arms

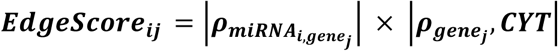

where 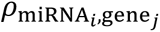 denotes the Spearman correlation coefficient between miRNA *i* and target gene *j*, and 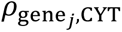 denotes the corresponding Spearman correlation coefficient between gene *j* and CYT.

The edge score was used exclusively to scale edge width in bipartite network visualisations and did not influence pair selection, recurrence assessment, machine-learning prioritisation, survival modelling, or any downstream analyses. No graph-theoretical analyses, including centrality estimation, hub detection, community detection, or topology inference, were performed.

### Tumor Purity Adjustment

To assess whether observed associations were confounded by variation in tumour cellularity, cross-cancer recurrent miRNA–gene–CYT triplets were re-evaluated after adjustment for tumour purity. Within each eligible cancer type, miRNA expression, target gene expression, and CYT values were independently regressed on tumour purity, and residuals from these models were used for subsequent analyses.

CPE was designated a priori as the primary tumour purity metric because it integrates multiple orthogonal purity estimators into a consensus measure. LUMP was included as an independent sensitivity metric to assess the robustness of recurrent associations to an alternative purity estimator (27).

For each purity metric, Spearman correlations were recomputed using purity-adjusted residuals, and BH correction was applied separately within each directional branch. Triplets were retained if they satisfied the predefined effect size and significance criteria (|ρ| ≥ 0.30 and FDR < 0.05) and demonstrated recurrence across at least three cancer types following purity adjustment.

Only triplets that remained significant after adjustment using the primary CPE metric were advanced to machine learning prioritization and survival analyses. Final pair selection and all downstream analyses were based exclusively on CPE-adjusted results, whereas LUMP-adjusted analyses were used solely for sensitivity assessment.

### Machine Learning: Random Forest and Bootstrap Stability

Candidate miRNAs derived from the CPE-adjusted recurrent pairs within each analysis track were evaluated for feature prioritisation using random forest regression (R package randomForest; random seed = 123) (28, 29). The continuous CYT score was used as the outcome variable. A full model comprising 1,000 decision trees was fitted for each track.

Feature importance was quantified using both node-purity increase (IncNodePurity) and permutation importance (%IncMSE). Bootstrap stability (30) was assessed across 100 resamples, with each resample fitted using a 300-tree random forest model. For each bootstrap iteration, the top 20 miRNAs ranked by IncNodePurity were recorded. Stability was defined as the proportion of bootstrap resamples in which a given miRNA appeared among the top 20 features. Model performance was evaluated using the out-of-bag (OOB) mean squared error (MSE) and OOB pseudo-R² obtained from the final random forest models.

A consensus machine-learning rank was computed as:

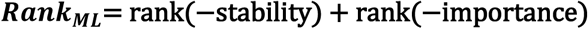

where lower values indicate higher overall priority.

Random forest modelling was performed exclusively after completion of all statistical filtering procedures and served solely as a feature-prioritization framework. Importantly, machine-learning outputs did not influence pair discovery, recurrence assessment, tumor-purity adjustment, or final pair selection. Final miRNA–gene pairs were defined exclusively by the CPE-adjusted cross-cancer recurrence analysis, whereas random forest analyses were used only to rank and prioritize the resulting candidate miRNAs for downstream interpretation.

### Survival Analysis

The prognostic significance of each candidate miRNA was evaluated using Cox proportional hazards regression models implemented in the R package survival (31, 32). OS time and event status were used as the outcome variables, and models were adjusted for cancer type to account for differences in baseline hazard across tumour entities. Cancer type was included as a categorical covariate to account for systematic differences across tumor types. For miRNA (*m*), the model was specified as:

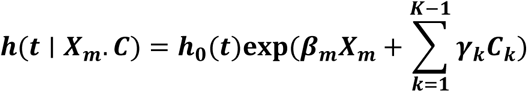

where (ℎ(*t* ∣ *X_m_*. *C*)) denotes the hazard function at time (*t*), (ℎ_0_(*t*)) is the baseline hazard function, (*X_m_*) represents the z-score-standardised expression value of miRNA (*m*), and (*C_k_*) denotes cancer-type indicator variables. The coefficient (*β_m_*) represents the log-hazard ratio (HRs) associated with a one-standard-deviation increase in miRNA expression.

Features exhibiting zero variance were excluded before model fitting. HR and 95% confidence intervals (CIs) were estimated for the miRNA expression term. BH correction was applied to miRNA-specific p-values within each analysis track. Models were fitted only when at least 50 complete observations spanning a minimum of two cancer types were available.

A composite miRNA signature score was constructed by averaging z-score-standardized expression values across the top 20 miRNAs ranked by the machine-learning consensus score and subsequently re-standardising the resulting score before Cox modelling. The final consensus prioritization score for miRNA (*i*) was defined as:

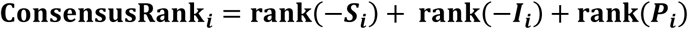

where (*S_i_*), (*I_i_*), and (*P_i_*) denote bootstrap stability, feature importance quantified by IncNodePurity, and the BH-adjusted Cox model p-value, respectively. Lower values of (ConsensusRank*_i_*) indicate higher overall priority.

### miRTarBase Experimental Validation Annotation

Following prioritization by machine-learning importance and survival significance, final miRNA–gene pairs were annotated against miRTarBase to assess experimental support. Two evidence sets were considered: all documented miRNA–target interactions and a subset restricted to strong experimental evidence, defined as reporter assay or western blot validation. Only human-to-human interactions (Homo sapiens miRNAs and target genes) were included, and miRNA identifiers were standardised to the hsa-prefix format prior to matching.

miRTarBase annotation was applied as a post hoc evidence layer and did not serve as a discovery filter; pairs lacking miRTarBase support were retained in the final recurrent miRNA–gene pair set.

### Mediation Analysis

To quantify the extent to which target genes mediated the association between miRNA expression and CYT, mediation analyses were performed for each miRNA–gene–CYT triplet using a regression-based framework (33) comprising sequential linear models and an analytical Sobel test (34). All models were fitted using the base R function *lm()*; no dedicated mediation package or bootstrap confidence interval were used. Cancer type was included as a categorical covariate in pan-cancer analyses involving more than one tumour type.

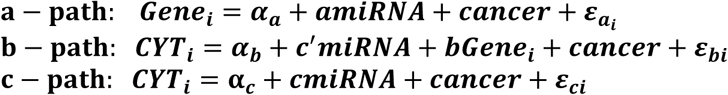

where (a) denotes the effect of miRNA expression on target gene expression, (b) denotes the effect of target gene expression on CYT after adjustment for miRNA expression, (c) denotes the total effect of miRNA expression on CYT, and (c’) denotes the direct effect after adjustment for target gene expression. Cancer type was included as a categorical covariate in all pan-cancer models

The indirect effect was estimated as:

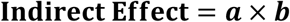

and the proportion mediated was calculated as:

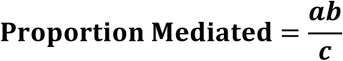

Statistical significance of the indirect effect was assessed using the Sobel test:

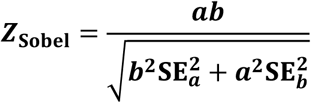

where 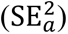 and 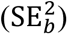 denote the standard errors of coefficients (*α*) and (*b*), respectively. Two-sided (p)-values were derived from the standard normal distribution, and BH correction was applied jointly across all pan-cancer Sobel tests, with significance defined as a FDR < 0.01.

Models were fitted only when at least 30 complete observations and a minimum of three distinct values per variable were available. Cancer-stratified mediation analyses were additionally performed within eligible tumour types, with BH correction applied independently within each cancer type.

Because mediation analyses were conducted using cross-sectional transcriptomic data, the results should not be interpreted as evidence of causality. Rather, they indicate statistical co-variation consistent with a gene-mediated association and should be considered hypothesis-generating findings requiring experimental validation.

### Immune Cell Integration

Immune cell abundance was estimated from mRNA expression data using a marker-gene scoring approach. Marker gene sets were defined using established cell-type-specific genes for eight immune cell populations: CD8⁺ T cells (CD8A, CD8B, CD3D, CD3E, CD247), NK cells (NCAM1, NKG7, KLRD1), regulatory T cells (FOXP3, IL2RA, IKZF2), B cells (CD19, MS4A1, CD79A), macrophages (CD68, CD163, MRC1), dendritic cells (ITGAX, CLEC9A, FLT3), neutrophils (FCGR3B, CEACAM8, CSF3R), and exhausted T cells (PDCD1, HAVCR2, LAG3, TIGIT, CTLA4).

For each marker gene present in the matched mRNA matrix, expression values were z-score standardized across all 9,288 samples. The per-sample score for each cell type was computed as the arithmetic mean of these gene-level z-scores. Cell types represented by fewer than two marker genes in the expression matrix were excluded. A ninth feature was computed as the pan-cancer z-score of CYT and served as an internal consistency reference. Associations between candidate miRNAs and all nine immune features were evaluated using Spearman correlation, both pan-cancer and within each eligible cancer type, with BH correction applied to cancer-stratified tests.

### Pan-Cancer Molecular Subtype Discovery

Pan-cancer molecular subtypes were identified by k-means clustering (35, 36) of miRNA expression profiles across all eligible tumor samples. The top *N* miRNAs from each regulatory track, ranked by the consensus framework, were selected as clustering features. Expression values were standardized across samples using z-score transformation.

K-means clustering was performed on the standardized expression matrix without batch-effect correction across a predefined range of cluster numbers ((k)), using 200 random initializations per (k). The optimal number of clusters was determined by consensus ranking of three metrics: mean silhouette width, the Calinski–Harabasz index, and the relative reduction in within-cluster sum of squares (elbow criterion). For each metric, candidate (k) values were ranked independently, and the consensus score was defined as the mean rank across metrics. Survival differences among molecular subtypes were evaluated using cancer-type-adjusted Cox proportional hazards models with subtype included as a categorical predictor. For visualization purposes only, cancer-type effects were removed from the standardized expression matrix prior to principal component analysis (PCA) (37). This correction was applied to low-dimensional visualization and did not affect subtype assignments, which were derived from the uncorrected standardized matrix.

### Pathway Enrichment Analysis

Target genes from the final pair rankings in each track were subjected to pathway enrichment analysis using clusterProfiler (38, 39). Gene Ontology (GO) (40) biological process and Kyoto Encyclopedia of Genes and Genomes (41) pathway terms were tested with the gene universe defined as all 20,311 genes present in the matched mRNA expression matrix. BH-corrected FDR < 0.05 was applied as the significance threshold for pathway enrichment. Enrichment results were computed separately for the negative and positive tracks.

### Drug–Target Analysis

Target genes from the final pair rankings across both tracks were queried against the DGIdb. Drug–target interaction hits were identified for each target gene and prioritized according to the miRNA consensus rank, to highlight targets most strongly supported by the multi-step miRNA discovery pipeline.

### Statistical Methods

All statistical analyses were performed in R. Spearman rank correlation was used for all pairwise association analyses. Multiple testing was controlled using the BH procedure throughout the study. An FDR threshold of 0.05 was applied for all discovery analyses, whereas mediation analyses used a more stringent FDR threshold of 0.01, as described in the corresponding methodological subsections. Random seeds were predefined to ensure reproducibility. A seed value of 123 was used for random forest modelling and bootstrap resampling, whereas a seed value of 42 was used for k-means clustering. All per-cancer analyses required a minimum of 30 matched samples per cancer type. Bootstrap confidence intervals were not used for mediation analyses; statistical significance was assessed exclusively using the analytical Sobel test.

## Results

### Pan-Cancer Cohort Characteristics and Discovery of 204 Recurrent miRNA Regulators of Cytolytic Activity

A total of 9,288 primary tumour samples with complete molecular and clinical information were retained for pan-cancer analyses, encompassing 31 TCGA cancer types (Supplementary Table S1). Cohort sizes varied substantially across tumour types, ranging from 36 cases in CHOL to 1,065 cases in BRCA (Figure 2A). Mature miRNA expression data were unavailable for GBM in the harmonized dataset; consequently, this tumour type was excluded from all downstream analyses. The final matched dataset comprised 743 mature miRNAs and 20,311 mRNA features.

**Figure 2.**
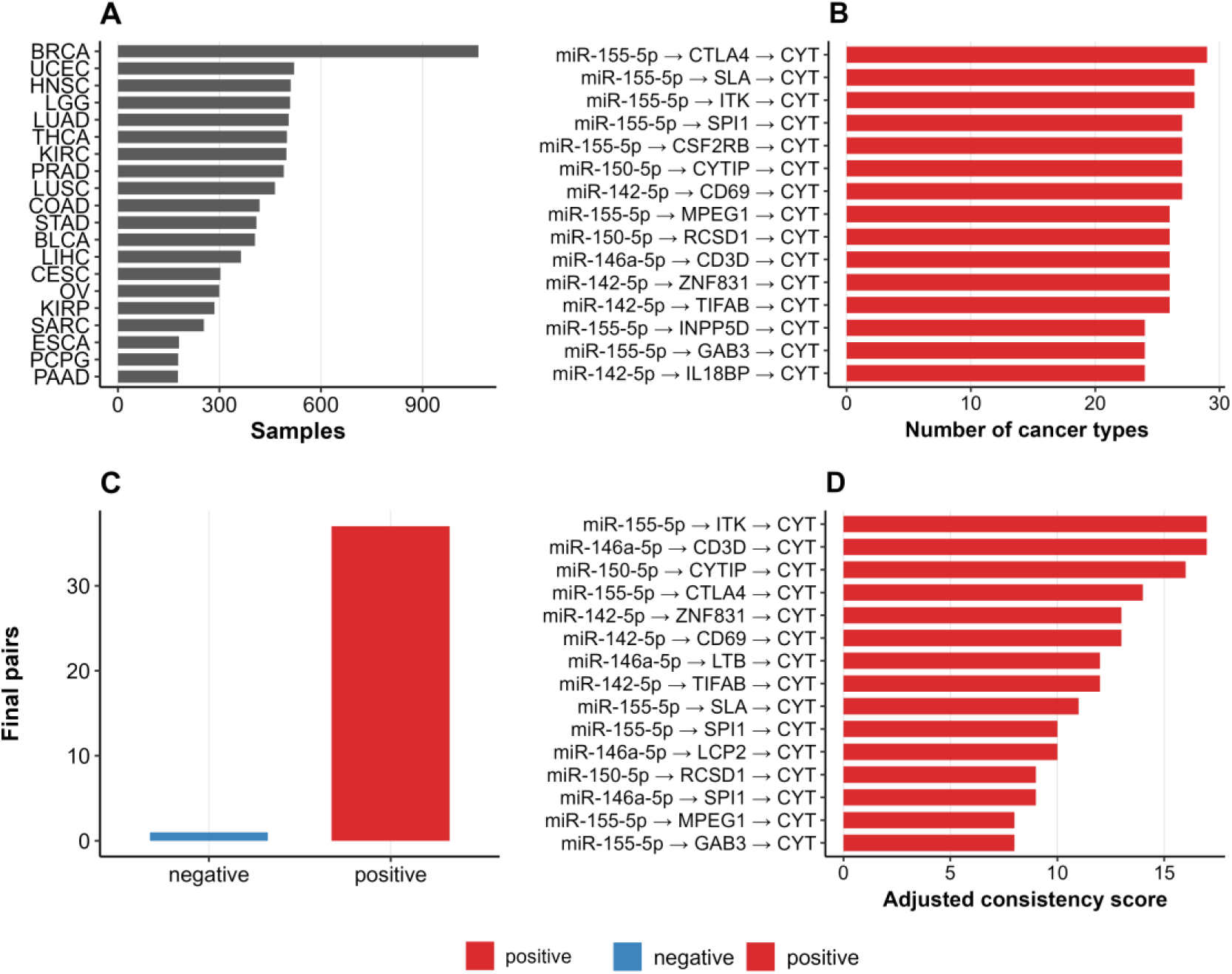
Pan-cancer cohort composition, cross-cancer recurrence, and prioritization of miRNA–gene– CYT associations. (A) Distribution of sample counts across cancer types included in the final matched analytical cohort following quality-control procedures. (B) Cross-cancer recurrence of the highest-ranking miRNA–gene–CYT triplets, quantified as the number of cancer types in which each triplet satisfied the predefined discovery criteria (Spearman |ρ| ≥ 0.30; FDR < 0.05). (C) Number of miRNAs–gene–CYT pairs retained within each regulatory track following recurrence filtering, tumour-purity adjustment, machine-learning prioritization, and survival-based selection. (D) Highest-ranking triplets according to the adjusted consistency score, a composite metric integrating cross-cancer recurrence and correlation strength. All triplets displayed in panels B and D originated from the positive regulatory track.

CYT scores were successfully derived for all 9,288 samples and demonstrated substantial heterogeneity across the pan-cancer cohort, with values ranging from 0 to 12.81 (mean = 6.50). OS data were available for 9,165 patients, among whom 2,493 death events were recorded. Cancer-specific correlation analyses between CYT scores and mature miRNA expression profiles generated 23,033 miRNA–cancer association tests. Following correction for multiple comparisons, 1,754 significant miRNA–CYT associations met the predefined significance criteria (FDR < 0.05 and |ρ| ≥ 0.30), involving 594 unique miRNAs across all 31 cancer types.

To identify reproducible immune-associated miRNAs, a cross-cancer recurrence analysis was performed, retaining only miRNAs that demonstrated significant associations with CYT in at least three independent tumour types. This approach yielded 204 candidate miRNAs for downstream integration analyses. The frequency of recurrence among these candidates ranged from three to 30 cancer types, highlighting marked differences in the consistency of miRNA–CYT relationships across malignancies. Among all candidates, hsa-miR-155-5p exhibited the highest degree of reproducibility, showing significant associations with CYT in 30 of the 31 analysed cancer types. Additional highly recurrent miRNAs included hsa-miR-155-5p and hsa-miR-142-5p, both of which displayed consistent positive associations with CYT across multiple tumour types (Fig. 2B). Application of the subsequent triple-filter integration framework identified a limited set of robust miRNAs–gene–CYT regulatory axes. The overwhelming majority of final associations were positive, whereas only a single negative association satisfied all selection criteria (Fig. 2C). Ranking of the integrated regulatory triplets using the adjusted consistency score further highlighted immune-related networks centred on hsa-miR-155-5p, hsa-miR-150-5p, hsa-miR-146a-5p, and hsa-miR-142-5p (Fig. 2D). The highest-scoring triplets included hsa-miR-155-5p–ITK–CYT, hsa-miR-146a-5p–CD3D–CYT, hsa-miR-150-5p–CYTIP–CYT, and hsa-miR-155-5p–CTLA4–CYT, all of which were recurrently observed across multiple tumour types and involved genes with established roles in T-cell activation and cytotoxic immune responses.

### Target Integration and Pan-Cancer Identification of 268 miRNA–Gene–CYT Regulatory Triplets

Target gene integration analyses were performed using the 204 recurrent CYT-associated miRNAs identified in the discovery phase. Predicted human miRNA–target interactions obtained from TargetScan were intersected with the matched mRNA expression matrix, resulting in 95 miRNAs with at least one predicted target gene represented in the dataset. Overall, this integration yielded 65,975 unique miRNA–gene pairs involving 20,311 expressed genes.

Pan-cancer correlation analyses across all 9,288 samples identified 268 miRNA–gene– CYT triplets that satisfied the predefined significance criteria, representing 50 unique miRNAs and 168 unique target genes. These retained associations reflected coordinated relationships between miRNA expression, target gene expression, and cytolytic activity across the pan-cancer cohort.

The distribution of significant triplets was markedly heterogeneous with respect to correlation directionality. Based on the signs of the miRNA–gene and gene–CYT associations, retained pairs were classified into four directional categories. The largest category comprised pairs exhibiting concordant positive associations between miRNA expression and target gene expression, as well as between target gene expression and CYT (154 pairs). In contrast, 77 pairs demonstrated negative miRNA–gene associations coupled with negative gene–CYT associations. A smaller subset of 22 pairs displayed positive miRNA–gene associations together with negative gene–CYT associations, whereas only 15 pairs showed negative miRNA–gene associations accompanied by positive gene–CYT associations (Supplementary Table S2).

When consolidated according to the direction of the miRNA–gene relationship, the positive regulatory track accounted for the majority of retained associations, comprising 176 miRNA–gene–CYT triplets (65.7%). The remaining 92 triplets (34.3%) belonged to the negative regulatory track. Overall, positive miRNA–gene associations were more frequent than negative associations among recurrent CYT-related miRNA–gene–CYT triplets identified across cancer types.

### Cross-Cancer Recurrence Analysis Identifies 149 Recurrent Triplets and Refines to 38 High-Confidence miRNA–Gene–CYT Regulatory Networks

Each of the 268 pan-cancer miRNA–gene–CYT triplets identified during the integration stage was re-evaluated across the 31 eligible cancer types to assess cross-cancer reproducibility. Application of the predefined recurrence criterion identified 149 recurrent triplets detected in at least three tumour types. These comprised 23 negative-track associations and 126 positive-track associations, indicating substantially greater cross-cancer reproducibility within the positive regulatory track.

In the negative regulatory track, 23 of 92 candidate triplets (25.0%) satisfied the recurrence threshold, involving 16 unique miRNAs and 19 target genes. Recurrence frequencies ranged from three to 13 cancer types. Among these associations, the hsa-miR-200b-3p– FLI1–CYT axis showed the highest degree of cross-cancer reproducibility, recurring in 13 tumour types. After downstream filtering, this association was retained as the sole final negative-track miRNA–gene–CYT triplet (Fig. 3).

**Figure 3.**
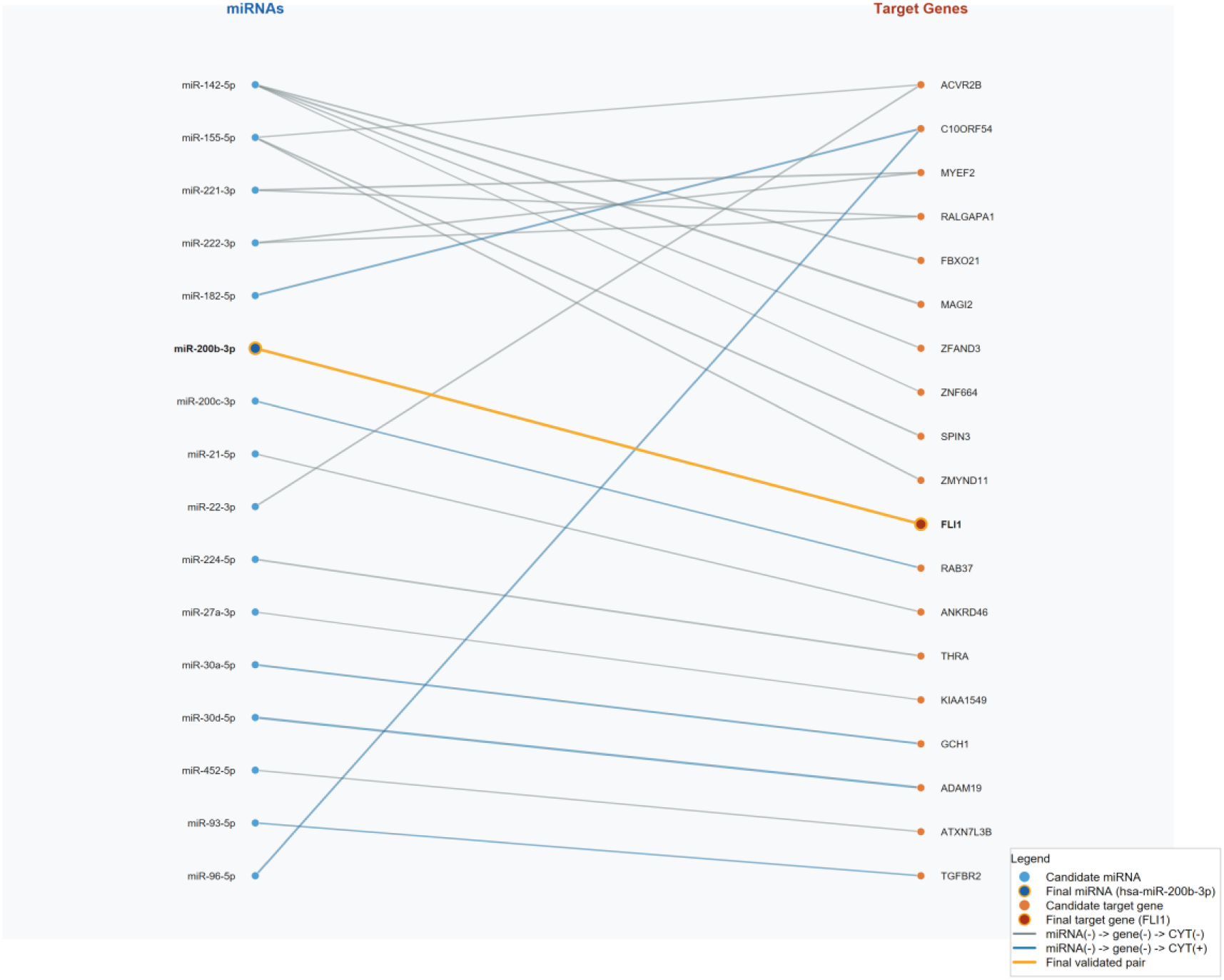
Network representation of recurrent negative-track miRNA–gene–CYT associations. Candidate miRNAs are shown on the left and predicted target genes on the right. Edges represent recurrent negative-track miRNA–gene–CYT triplets that satisfied the predefined recurrence criterion (detection in ≥3 cancer types). The highlighted hsa-miR-200b-3p–FLI1 association corresponds to the most recurrent negative-track triplet identified across tumour types and was retained as the final negative-track candidate for downstream analyses.

In contrast, 126 of 176 positive-track candidate triplets (71.6%) satisfied the recurrence threshold, involving 26 unique miRNAs and 95 target genes. Individual positive-track triplets were detected in three to 29 cancer types, with the hsa-miR-155-5p–CTLA4–CYT axis representing the most recurrent association. Following tumour-purity adjustment, machine-learning prioritization, and survival evaluation, 37 positive-track triplets involving eight miRNAs and 30 target genes were retained in the final validated positive-track network (Supplementary Figure S1).

Overall, sequential filtering reduced the 149 recurrent triplets to 38 high-confidence miRNA–gene–CYT triplets, comprising one negative-track association and 37 positive-track associations. These final candidates involved nine unique miRNAs and 31 unique target genes. Detailed recurrence counts, tumour types contributing to each retained association, random forest metrics, survival estimates, and ranking information are provided in Supplementary Table S3.

### Tumor Purity Adjustment, Prioritization, and Prognostic Evaluation

The 149 cross-cancer recurrent miRNA–gene–CYT triplets (23 negative-track and 126 positive-track associations) were evaluated for robustness against tumor purity. Within each cancer type, miRNA expression, target gene expression, and CYT values were residualized against tumor purity, and associations were re-assessed using the original correlation and significance criteria.

CPE as the primary metric, 38 triplets remained significant after adjustment, comprising one negative-track association and 37 positive-track associations (Fig. 2C; Supplementary Table S3). The sole negative-track triplet retained was hsa-miR-200b-3p–FLI1–CYT (recurrent in three cancer types), while the positive-track triplets involved eight unique miRNAs and 30 unique target genes. Sensitivity analyses using LUMP confirmed greater robustness in the positive regulatory track (66 recurrent triplets).

Candidate miRNAs were further prioritized using random forest regression with CYT as the outcome variable. The positive-track model demonstrated better predictive performance than the negative-track model, with a lower out-of-bag (OOB) mean squared error (1.158 vs. 2.869) and a higher OOB pseudo-R² (0.694 vs. 0.243), whereas both models were trained using identical bootstrap and random forest settings (Supplementary Table S4). Among the 19 candidate miRNAs evaluated by random forest, all nine miRNAs retained in the final validated network demonstrated complete bootstrap stability (stability = 1.00) across 100 resampling iterations (Figs. 4B and 4E; Supplementary Table S5). hsa-miR-200b-3p showed the highest feature importance in the negative track, whereas hsa-miR-155-5p ranked highest in the positive track (Figs. 4A and 4D).

**Figure 4.**
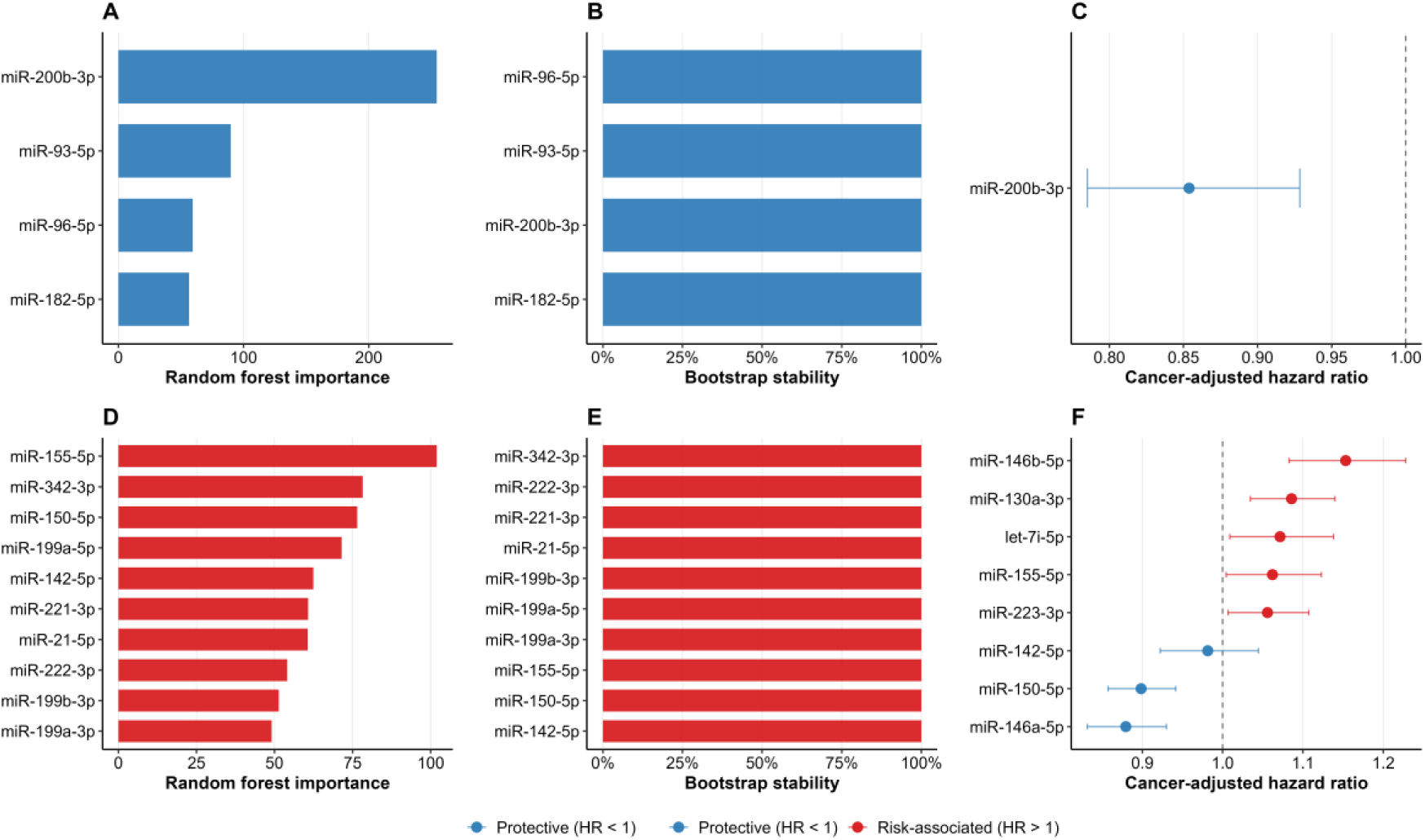
Prioritization and survival evaluation of recurrent pan-cancer miRNA candidates after tumour-purity adjustment. **(A)** Random forest feature-importance ranking of negative-track miRNAs identified from the LUMP-adjusted recurrent candidate set. hsa-miR-200b-3p exhibited the highest predictive importance among negative-track candidates. **(B)** Bootstrap stability analysis of negative-track miRNAs, showing the proportion of resampling iterations in which each candidate was retained. hsa-miR-96-5p, hsa-miR-93-5p, hsa-miR-200b-3p, and hsa-miR-182-5p achieved complete stability across bootstrap iterations. **(C)** Cancer-type-adjusted Cox proportional hazards analysis for the final negative-track candidate, hsa-miR-200b-3p, demonstrating a protective association with OS (HR < 1). **(D)** Random forest feature-importance ranking of positive-track miRNAs derived from the LUMP-adjusted recurrent candidate set. hsa-miR-155-5p showed the greatest predictive importance, followed by hsa-miR-342-3p and hsa-miR-150-5p. **(E)** Bootstrap stability analysis of positive-track miRNAs, illustrating highly reproducible candidate selection across resampling iterations. **(F)** Cancer-type-adjusted Cox proportional hazards analysis of the final positive-track miRNA candidates. Error bars represent 95% CI. Blue markers indicate protective associations (HR < 1), whereas red markers indicate risk-associated relationships (HR > 1). HR were estimated using pan-cancer Cox models adjusted for cancer type. Only miRNAs satisfying recurrence, tumour-purity adjustment, machine-learning prioritization, and survival-selection criteria were retained for downstream analyses.

Subsequent cancer-type-adjusted Cox proportional hazards modeling demonstrated that eight of the nine miRNAs had independent prognostic value. hsa-miR-200b-3p was associated with improved survival in the negative track (HR = 0.854, 95% CI 0.785–0.929, FDR = 2.23 × 10⁻⁴). In the positive track, hsa-miR-146a-5p and hsa-miR-150-5p showed protective effects, whereas hsa-miR-146b-5p, hsa-miR-130a-3p, hsa-miR-155-5p, hsa-let-7i-5p, and hsa-miR-223-3p were associated with increased risk (Figs. 4C and 4F; Supplementary Figure S2).

### Final Selection of High-Confidence miRNA–Gene–CYT Triplets

After integrating cross-cancer recurrence, CPE-based tumor purity adjustment, machine-learning prioritization, and survival analysis, 38 high-confidence miRNA–gene–CYT triplets were retained. These comprised one negative-track association (hsa-miR-200b-3p– FLI1–CYT) and 37 positive-track associations involving a total of nine unique miRNAs and 31 unique target genes (Supplementary Table S3). This final set formed the basis for all subsequent mechanistic and clinical analyses.

### Mediation Analysis and Immune Cell Associations

Pan-cancer mediation analysis was performed for all 38 triplets using cancer-type-adjusted linear models and the Sobel test. Significant gene-level mediation was confirmed for 37 triplets (FDR < 0.01), with the proportion mediated ranging from 0.08 to 0.94 (Supplementary Table S6). The strongest mediation effects were observed for hsa-miR-200b-3p–FLI1–CYT (94.2%) and hsa-miR-146a-5p–CD3D–CYT (93.7%).

Immune cell integration further supported the biological relevance of the signature. Of 171 tested associations, 161 remained significant pan-cancer (FDR < 0.05). The composite miRNA signature score correlated strongly with multiple immune populations, particularly Treg cells (ρ = 0.359), exhausted T cells (ρ = 0.335), B cells (ρ = 0.334), and CD8⁺ T cells (ρ = 0.333) (Fig. 5C). Positive-track miRNAs consistently showed stronger immune associations than negative-track candidates (Fig. 5D), highlighting predominant links to adaptive immune activation in the TME (Figs. 5A and 5B).

**Figure 5.**
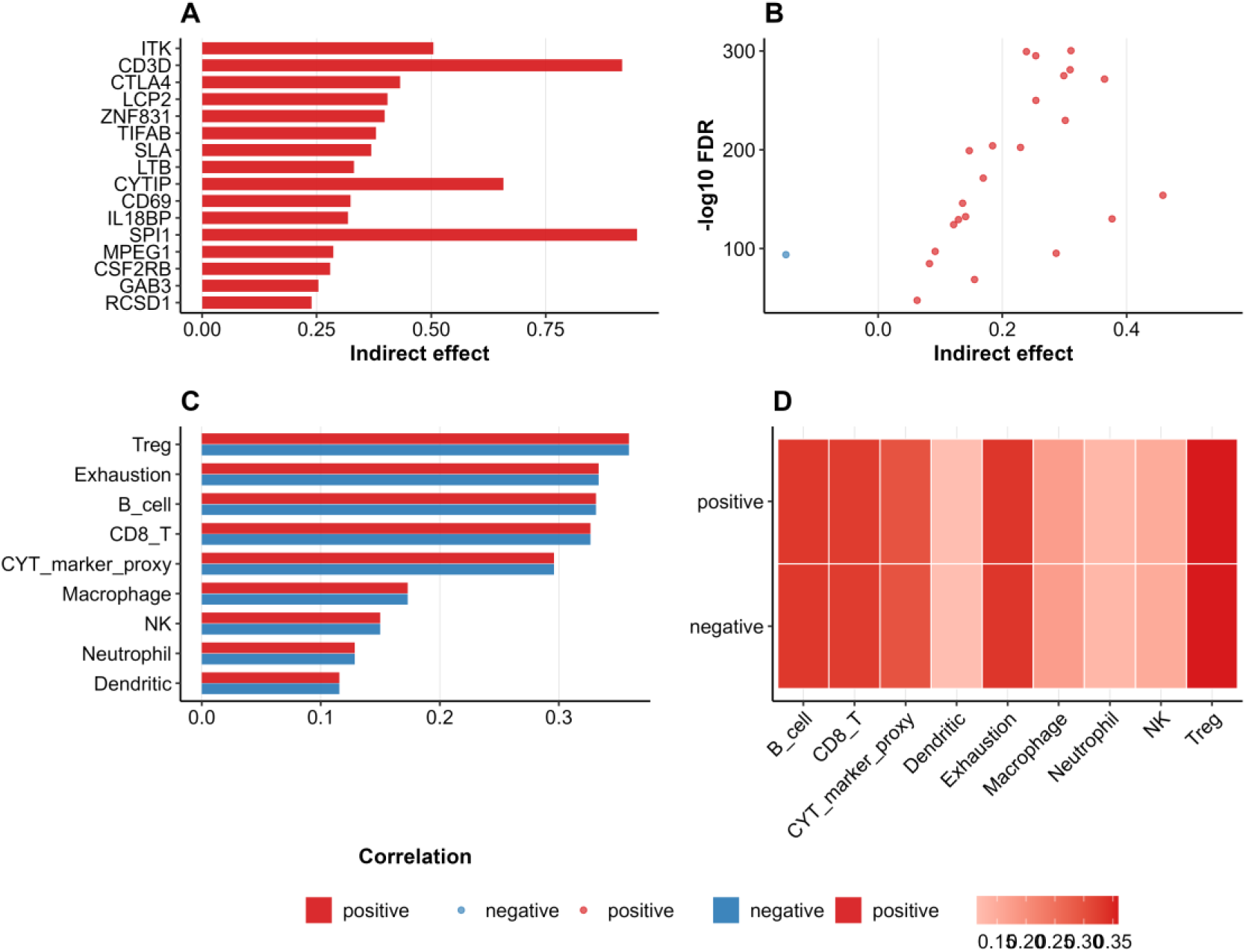
Integration of mediation and immune-cell association analyses. **(A)** Indirect mediation effects for the top mediator genes identified among the final miRNA–gene–CYT triplets. **(B)** Relationship between mediation effect size and statistical significance −log10 (FDR) across significant mediated associations. **(C)** Spearman correlations between the composite miRNA signature score and nine immune-related features, including B cells, CD8⁺ T cells, regulatory T cells (Treg), exhausted T cells, macrophages, dendritic cells, neutrophils, NK cells, and a CYT-derived reference score. **(D)** Heatmap summarizing immune-feature correlations for positive- and negative-track candidate miRNAs. Positive-track candidates showed consistently stronger associations with immune-cell infiltration across most immune populations, particularly lymphocyte-related features, supporting their close relationship with tumour immune activation.

### Pan-Cancer Molecular Subtype Discovery

To assess whether the final miRNA signature could define clinically meaningful tumor subgroups, unsupervised k-means clustering was performed using the 19 non-redundant candidate miRNAs across 9,288 samples. A two-cluster solution was optimal based on silhouette width, Calinski–Harabasz index, and elbow criterion (Fig. 6D).

**Figure 6.**
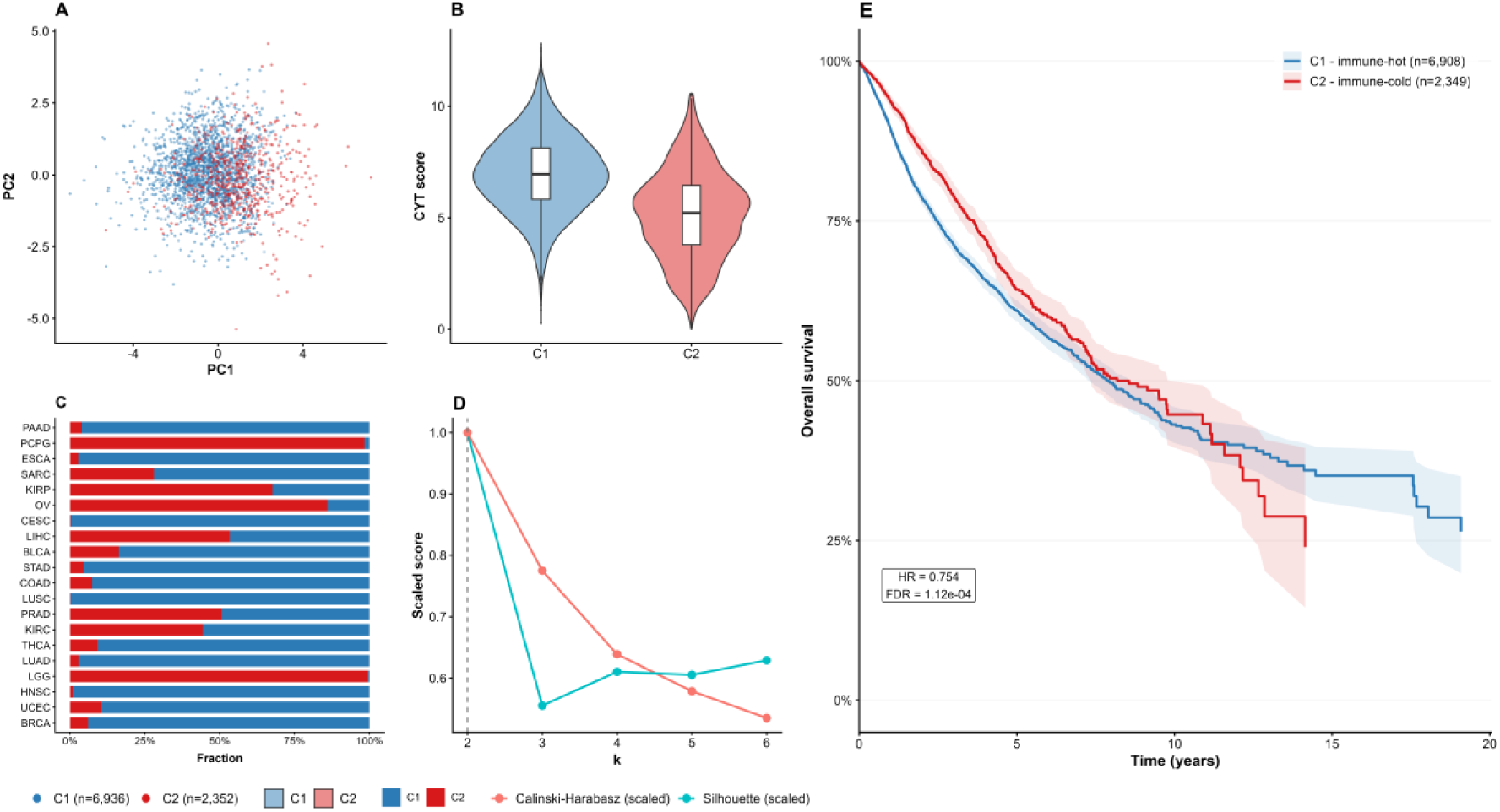
Pan-cancer molecular subtype discovery based on the final miRNA signature. **(A)** PCA of the 19-miRNA expression signature across 9,288 TCGA tumour samples, showing separation of the two unsupervised molecular subtypes identified by k-means clustering. **(B)** Distribution of CYT scores in the two subtypes. The C1 subtype exhibited significantly higher CYT scores than C2 and was therefore designated the immune-hot subtype, whereas C2 was classified as immune-cold. **(C)** Distribution of the two molecular subtypes across individual cancer types, illustrating differences in subtype composition among tumour cohorts. **(D)** Evaluation of clustering performance for *k* = 2–6 using the scaled Calinski–Harabasz index and silhouette coefficient. Both metrics supported a two-cluster solution, which was selected for downstream analyses. **(E)** Kaplan–Meier OS curves for the immune-hot (C1) and immune-cold (C2) subtypes in the cancer-type-adjusted survival cohort. Shaded regions represent 95% CIs. HR and FDR were estimated using a cancer-type-adjusted Cox proportional hazards model.

The two subtypes (C1: n=6,936; C2: n=2,352) were designated as immune-hot (C1) and immune-cold (C2) based on marked differences in CYT scores (Fig. 6B). Both subtypes were distributed across multiple cancer types (Fig. 6C), and PCA confirmed clear transcriptomic separation (Fig. 6A). Cancer-type-adjusted survival analysis demonstrated significantly better OS in the immune-hot subtype (HR = 0.754, FDR = 1.12 × 10⁻⁴; Fig. 6E).

### Functional Enrichment, Network Topology, and Drug-Target Prioritization of the miRNA–Gene–CYT Regulatory Network

Functional enrichment analysis was performed on the 30 unique target genes from the positive regulatory track (the dominant track in the final network), using all 20,311 expressed genes as the background. Significant enrichment (FDR < 0.05) was observed for multiple immune-related GO biological processes, most notably lymphocyte differentiation, T-cell differentiation, regulation of leukocyte activation, antigen receptor-mediated signaling, and T-cell receptor signaling (Fig. 7A; Supplementary Table S7). These findings confirm strong enrichment of the miRNA–gene–CYT network for adaptive immune processes and immune-cell activation pathways. Network topology analysis revealed several hub genes regulated by multiple candidate miRNAs. SPI1 emerged as the most highly connected hub, followed by SOCS1, LCP2, CYTIP, CD80, and CD3D (Fig. 7B). The central positioning of these genes suggests they serve as key intermediaries linking miRNA regulation to tumor immune activity.

**Figure 7.**
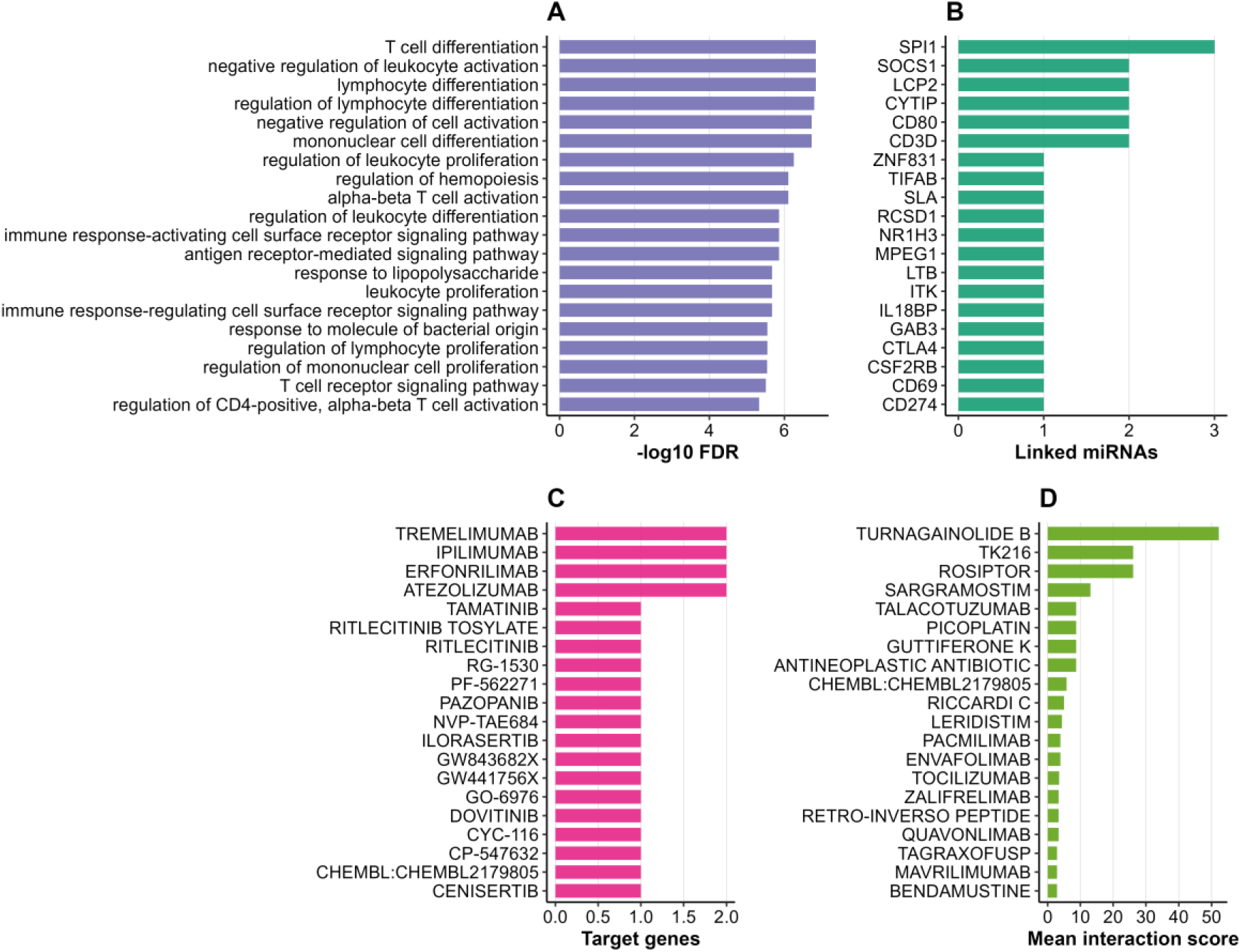
Functional enrichment and drug–target prioritization of the final miRNA–gene–CYT network. **(A)** Top significantly enriched GO biological processes identified from the final target-gene set, ranked by −log10(FDR). Enriched categories were predominantly related to lymphocyte differentiation, T-cell activation, leukocyte regulation, and immune receptor signalling. **(B)** Network connectivity analysis showing the number of candidate miRNAs linked to each target gene. SPI1 was the most highly connected hub gene, followed by SOCS1, LCP2, CYTIP, CD80, and CD3D. **(C)** Top therapeutic compounds ranked according to the number of target genes represented within the final network. Immune checkpoint inhibitors, including tremelimumab, ipilimumab, and atezolizumab, were among the highest-ranking candidates. **(D)** Top compounds ranked by mean drug–target interaction score, highlighting agents with the strongest aggregate evidence across network targets. These results demonstrate the enrichment of immune-related pathways and identify multiple pharmacologically actionable targets within the final miRNA–gene–CYT network.

To assess translational potential, drug–gene interactions were queried for all 31 target genes using the DGIdb database. Eleven target genes were identified as druggable, yielding 124 unique drug–gene interaction entries corresponding to 124 unique compounds (Supplementary Table S8). These included clinically relevant immune checkpoint genes such as CTLA4 and CD274 (PD-L1). When prioritized by the number of supported targets, prominent compounds included the immune checkpoint inhibitors tremelimumab, ipilimumab, and atezolizumab (Fig. 7C). Alternative ranking by mean interaction score highlighted additional agents targeting immune-regulatory pathways, such as turnagainolide B, TK216, and rosiptor (Fig. 7D). Collectively, these analyses demonstrate that the final miRNA–gene–CYT network is highly enriched for adaptive immune signaling pathways and contains multiple pharmacologically actionable targets. This supports the biological relevance of the identified regulators and highlights their potential for biomarker development, patient stratification, and drug repurposing in cancer immunotherapy.

## Discussion

In this study, we performed a comprehensive pan-cancer analysis of 9,288 primary tumours across 31 cancer types. We identified 38 recurrent miRNA–gene–CYT triplets involving nine conserved miRNAs and 31 target genes. These associations remained robust after stringent tumour purity adjustment, cross-cancer recurrence filtering, machine-learning prioritization with bootstrap stability assessment, and formal mediation testing. Together, these steps established a high-confidence regulatory network linking miRNA expression to cytolytic anti-tumour immunity.

A prominent feature of the network was the marked asymmetry between regulatory tracks, with 37 of the 38 final triplets belonging to the positive regulatory track and only one negative-track association (hsa-miR-200b-3p–FLI1–CYT) surviving all filtering steps. This predominance likely reflects both biological reality and the inherent characteristics of bulk tumor transcriptomics. Coordinated upregulation of miRNAs and target genes within infiltrating immune-cell populations can produce concordant positive correlations, even when canonical post-transcriptional repression is operative at the single-cell level.

Mediation analysis provided strong statistical support for gene-mediated mechanisms. Significant indirect effects were confirmed in 37 of the 38 triplets, with the proportion mediated reaching up to 94%. The strongest mediation effects were observed for hsa-miR-*200b-3p–FLI1–CYT* and *hsa-miR-146a-5p–CD3D–CYT*. hsa-miR-200b-3p was the only miRNA retained in the negative regulatory track. Although the miR-200 family is classically associated with inhibition of epithelial to mesenchymal transition, FLI1 is a key haematopoietic transcription factor involved in immune cell development and differentiation. The strong mediation effect in this axis suggests that hsa-miR-200b-3p modulates cytolytic activity through broader effects on immune cell composition and transcriptional programming in the TME (42, 43). hsa-miR-146a-5p exhibited extensive cross-cancer reproducibility and robust mediation via CD3D, a central component of T-cell receptor signaling (44). Although canonically viewed as a negative regulator of Nuclear Factor kappa-light-chain-enhancer of activated B cells driven inflammation (45, 46), its positive association with cytolytic activity here indicates highly context-dependent functions that are shaped by immune cell composition and activation status within the TME (47). Also, hsa-miR-155-5p emerged as a central hub miRNA in the network. It was recurrently associated with multiple immune-related targets, including CTLA4, ITK, CD274 (PD-L1), SPI1, and SOCS1. These findings extend previous work establishing miR-155 as a master regulator of innate and adaptive immunity, including macrophage activation, T-cell differentiation, and B-cell function (8, 12, 13). Our results demonstrate its broad involvement in coordinating tumor-associated immune activation and cytolytic activity across diverse cancer types.

The remaining miRNAs in the final signature (hsa-miR-150-5p, hsa-miR-142-5p, hsa-miR-146b-5p, hsa-let-7i-5p, and hsa-miR-223-3p) also exhibited consistent associations with immune-related genes and pathways. These miRNAs have well-documented roles in immune cell biology. For instance, hsa-miR-150-5p is a key regulator of B-cell and T-cell differentiation and maturation (48, 49). hsa-miR-142-5p participates in hematopoietic lineage specification and modulates both innate and adaptive immune responses (50). hsa-miR-146b-5p, like its paralog miR-146a, negatively regulates inflammatory signaling while showing context-dependent effects in tumor-infiltrating lymphocytes (51). Members of the let-7 family, including hsa-let-7i-5p, control T-cell activation and cytokine production (52), whereas hsa-miR-223-3p is a critical regulator of myeloid cell differentiation, neutrophil function, and macrophage polarization (53). All nine miRNAs demonstrated complete bootstrap stability and showed pervasive correlations with multiple immune cell populations, most notably B cells, CD8⁺ T cells, regulatory T cells, and exhausted T cells. Collectively, these miRNAs appear to orchestrate coordinated rather than isolated immune regulatory programmes within the TME. This coordinated action likely contributes to the ability of the nine-miRNA signature to define clinically meaningful pan-cancer immune subtypes.

Functional enrichment and network topology analyses further reinforced the immune-centric nature of the identified regulatory program. Target genes from the final network were strongly enriched for GO biological processes related to lymphocyte differentiation, T-cell activation, antigen receptor signaling, and leukocyte regulation. These findings align with the central role of the miRNA signature in modulating adaptive anti-tumor immunity. Network topology analysis revealed several highly connected hub genes, including SPI1, SOCS1, LCP2, CYTIP, CD80, and CD3D. These hubs likely serve as critical integration nodes through which multiple miRNAs converge to fine-tune immune responses within the TME. SPI1 (PU.1) is a master transcription factor essential for myeloid and lymphoid cell development (54). SOCS1 is a key negative regulator of cytokine signaling that modulates T-cell activation and prevents excessive inflammation (55, 56). LCP2 (SLP-76) is an adaptor protein critical for T-cell receptor signalling and cytotoxic function (57). CYTIP regulates cytoskeletal dynamics and T-cell adhesion (58), while CD80 and CD3D are central to co-stimulatory signaling and T-cell receptor complex assembly, respectively (44, 59). The convergence of multiple miRNAs on these hubs underscores their importance as downstream effectors in the regulation of anti-tumor cytolytic immunity.

Unsupervised k-means clustering based on the nine-miRNA signature identified two biologically distinct pan-cancer immune subtypes that transcend conventional tumor histology. We designated these as immune-hot (C1) and immune-cold (C2) based on marked differences in cytolytic activity and immune cell infiltration. The immune-hot subtype exhibited significantly higher CYT scores, greater infiltration of CD8⁺ T cells, B cells, and other effector populations, and was associated with substantially better OS after cancer-type adjustment (HR = 0.754, FDR = 1.12 × 10⁻⁴). In contrast, the immune-cold subtype showed lower cytolytic activity and poorer clinical outcomes. This miRNA-driven classification aligns closely with the established immune-hot and immune-cold tumor paradigm in immuno-oncology (60). Notably, several miRNAs in our signature have been previously implicated in shaping these phenotypes. In particular, hsa-miR-155-5p, a central hub in our network, has been consistently associated with immune-hot tumors and enhanced anti-tumor immunity in multiple studies (12, 13, 61). Elevated miR-155 expression promotes pro-inflammatory signaling, T-cell activation, and increased lymphocyte infiltration, features characteristic of immune-hot microenvironments (12, 14, 61). Similarly, hsa-miR-146a-5p and hsa-miR-142-5p have been linked to modulation of tumor inflammation and immune cell recruitment (45, 50, 62). These prior observations support our finding that the nine-miRNA signature captures shared regulatory programs driving immune phenotypes across cancer types, independent of tissue-of-origin. Such miRNA-based classifiers may therefore offer complementary biomarkers for predicting immunotherapy response and refining patient stratification strategies.

From a translational perspective, the identification of clinically actionable targets such as CTLA4 and CD274 (PD-L1) within the miRNA–gene–CYT network highlights the therapeutic potential of this signature. Several hub genes regulated by the nine miRNAs encode proteins that are already established targets of approved immunotherapies. For example, CTLA4 and PD-L1 (encoded by CD274) are the primary targets of ipilimumab, tremelimumab, and various PD-1/PD-L1 inhibitors that have transformed the treatment landscape of multiple cancers. The fact that these genes are under the regulatory influence of our conserved miRNAs suggests that the signature may not only reflect the immune state of the tumor but could also help predict or modulate responsiveness to checkpoint blockade (63). In addition to these well-known checkpoints, DGIdb analysis nominated 124 candidate compounds targeting 11 druggable genes in the network. These include both approved agents and investigational compounds with immune-modulatory properties. This substantial druggability potential positions the identified miRNA network as a valuable resource for drug repurposing and combination therapy strategies in immuno-oncology. Future studies integrating this miRNA signature with clinical trial data may help identify patients most likely to benefit from specific immunotherapeutic regimens and guide the development of miRNA-based therapeutics or mimics to reshape the tumor immune microenvironment.

### Strengths and Limitations

This study has several notable strengths, including the large pan-cancer cohort encompassing 31 tumor types, the integration of multiple molecular layers, rigorous tumour-purity adjustment, and the requirement for cross-cancer reproducibility. The use of complementary analytical approaches, including machine learning, survival modelling, mediation analysis, immune-cell integration, pathway enrichment, molecular subtyping, and druggability assessment provided convergent evidence supporting the robustness of the final miRNA–gene–CYT network.

Several limitations should also be acknowledged. First, all analyses were based on retrospective observational TCGA data and therefore cannot establish causal relationships. Second, external validation in independent cohorts was not performed. Such validation remains challenging because few publicly available datasets concurrently provide matched miRNA expression, mRNA expression, tumor-purity estimates, and long-term clinical outcome data across multiple cancer types. Third, target-gene assignments were based primarily on computational prediction and therefore require experimental confirmation. Fourth, CYT represents a transcriptomic surrogate of cytolytic immune activity rather than a direct measurement of immune-cell killing function. Fifth, the use of bulk transcriptomic data precludes cell-type-specific resolution and may contribute to the predominance of positive miRNA–gene associations observed in the final network. An additional limitation arises from the preprocessing strategy required for pan-cancer harmonization. Following batch-effect correction and quality-control procedures, 743 miRNAs were retained for downstream analyses. Consequently, a substantial number of miRNAs present in the original TCGA datasets were excluded because of missing values, low coverage, or incompatibility across tumor types. Although this filtering step improved cross-cancer comparability and reduced technical variation, it may also have resulted in the omission of biologically relevant immune-associated miRNAs that could not be robustly analysed within the unified pan-cancer framework. Accordingly, the identified miRNA–gene–CYT triplets should be regarded as high-confidence, hypothesis-generating candidates rather than experimentally validated regulatory relationships.

### Future Directions

Future research should prioritize functional validation of the highest-confidence regulatory triplets using CRISPR-based editing, reporter assays, and immune-cell co-culture models. Single-cell and spatial transcriptomics will be instrumental in dissecting cell-type-specific contributions. Prospective validation of the miRNA-derived immune subtypes in immunotherapy-treated cohorts will clarify their predictive utility. Finally, therapeutic modulation of these miRNAs or their downstream targets may open new avenues for enhancing anti-tumour immunity.

## Conclusions

This study presents a comprehensive, tumor purity-adjusted pan-cancer framework that systematically maps miRNA-associated regulatory networks governing cytolytic anti-tumor immunity across 31 human cancer types from TCGA. By integrating multi-layered analyses, we identified 38 high-confidence miRNA–gene–CYT regulatory triplets involving nine conserved miRNAs and 31 target genes. These associations exhibited strong reproducibility, with nearly all triplets showing significant gene-mediated effects and complete bootstrap stability across resamples. The resulting nine-miRNA signature defined two distinct pan-cancer immune subtypes (immune-hot versus immune-cold) that transcend conventional cancer-type boundaries, displaying marked differences in cytolytic activity and OS. Functional characterization revealed enrichment for T-cell activation, lymphocyte differentiation, and antigen receptor signaling pathways, while drug-target analysis highlighted multiple clinically actionable targets, including CTLA4 and CD274 (PD-L1), nominating 124 candidate therapeutic compounds. Collectively, these findings showed a compact and highly reproducible set of miRNA regulators of cytolytic immunity, provide biomarkers for immune subtyping, and reveal actionable nodes within the tumor immune microenvironment. This study offers a valuable resource for biomarker development, patient stratification, and future immuno-oncology research aimed at enhancing cancer immunotherapy strategies.

## Declarations

### Author Contributions

**N.B. (Nazanin Bagherlou):** Conceptualization, Methodology, Software, Formal analysis, Data curation, Visualization, Writing – original draft. **S.A. (Shahram Aliyari):** Supervision, Conceptualization, Methodology, Project administration, Writing – review & editing. **Z.S. (Zahra Salehi):** Resources, Data curation, Writing – review & editing. **M.P. (Mobin Pirouzkhah):** Validation, Writing – review & editing. **C.-A.W. (Cleo-Aron Weis):** Senior supervision, Methodology, Validation, Investigation, Writing – review & editing. All authors have read and agreed to the published version of the manuscript.

### Data Availability

All primary data used in this study are publicly available. Pan-cancer miRNA expression and mRNA expression data were obtained from the UCSC Xena TCGA Pan-Cancer Atlas hub (https://xenabrowser.net/; accessed January 2024) (19). Survival and clinical annotation data were obtained from Liu et al. (20). CYT scores were derived from GZMA and PRF1 expression as defined by Rooney et al. (1) and Thorsson et al. (2). TargetScan v7 predictions (22) were used for miRNA–target gene integration. All data are available at the sources cited; no new data were generated.

### Code Availability

The source code for the Pan-Cancer miRNA–Gene–CYT Analysis Pipeline is publicly available on GitHub at https://github.com/BagherlouBioinfo/PanCancer-miRNA-Gene-CYT, and has been permanently archived on Zenodo. The exact version used for all analyses presented in this study is version v1.0.0 and is accessible via https://doi.org/10.5281/zenodo.20999443

## Acknowledgements

The authors acknowledge TCGA Research Network and the UCSC Xena team for providing open-access pan-cancer genomic data. The authors thank the developers of the randomForest, clusterProfiler, survival, TargetScan, and DGIdb tools for open-access software and resources.

## Ethics Statement

This study used publicly available, de-identified data from TCGA, collected under existing institutional review board governance and patient informed consent procedures. No new biological samples were collected or analysed, and no additional ethics approval was required.

## Supplementary Materials Statement

**Supplementary Materials:** The following supporting information can be downloaded at the journal’s website:

**Table S1**, cancer-type composition of the final analytical cohort;

**Table S2**, recurrent miRNA–gene–CYT triplets identified before tumour-purity adjustment;

**Table S3**, final CPE-adjusted miRNA–gene–CYT triplets retained after integrated prioritization;

**Table S4**, random forest feature-prioritization results for candidate miRNAs;

**Table S5**, random forest model performance metrics;

**Table S6**, pan-cancer mediation analysis results;

**Table S7**, functional enrichment analysis of target genes from the final miRNA–gene– CYT network;

**Table S8**, drug–target prioritization results derived from DGIdb.

All supplementary tables are provided in a single file: **Supplementary_Tables.xlsx**.

## Abbreviations

BH: Benjamini–Hochberg multiple testing correction
CIs: Confidence Intervals
CPE: Consensus Purity Estimate
CYT: Cytolytic Activity
DGIdb: Drug Gene Interaction Database
EMT: Epithelial-to-Mesenchymal Transition
EB-adjusted: Bayes-adjusted
FDR: False Discovery Rate
GO: Gene Ontology
GZMA: Granzyme A
HR: Hazard Ratio
KEGG: Kyoto Encyclopedia of Genes and Genomes
KM: Kaplan–Meier
LUMP: Leukocytes Unmethylation for Purity
miRNA: microRNA
NF-κB: Nuclear Factor kappa-light-chain-enhancer of activated B cells
OOB: Out-of-Bag (random forest error rate)
OS: Overall Survival
PCA: Principal Component Analysis
PRF1: Perforin 1
RF: Random Forest
TCGA: The Cancer Genome Atlas
TME: Tumour Microenvironment
UMAP: Uniform Manifold Approximation and Projection

## Declaration of generative AI and AI-assisted technologies in the writing process

During the preparation of this work, the author(s) used ChatGPT for language editing, improving readability, grammar refinement, and rephrasing of sentences. After using this tool/service, the author(s) reviewed and edited the output as needed and take full responsibility for the content of the published article.

