## Supplementary_Figure S1,S2 for "Pan-cancer analysis identifies nine conserved miRNA regulators of tumor cytolytic activity and clinically actionable immune targets"

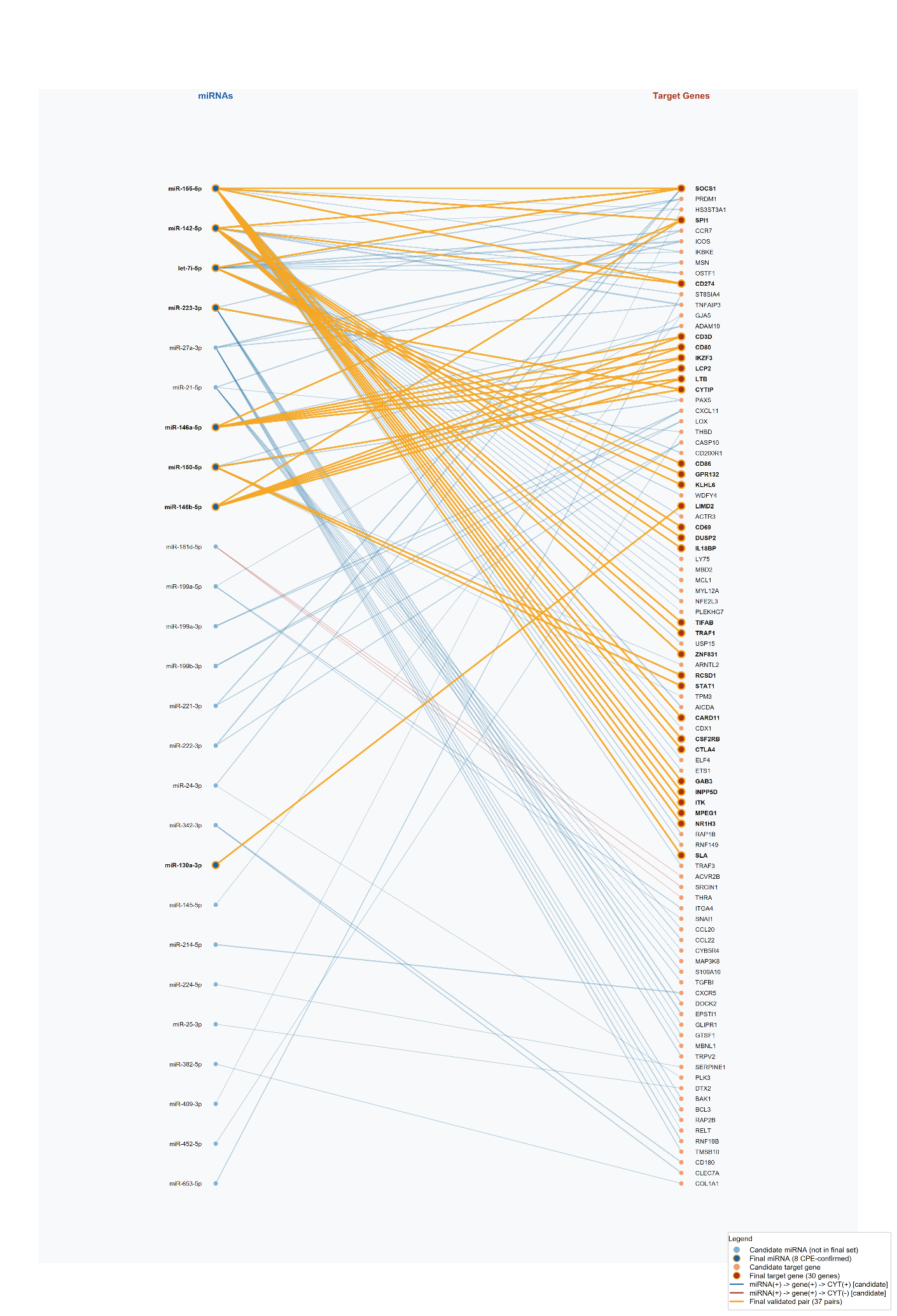

**Supplementary Figure S1. Network representation of recurrent positive-track miRNA–gene–CYT associations.** Candidate miRNAs are shown on the left and predicted target genes on the right. Edges represent recurrent positive-track miRNA–gene–CYT triplets that satisfied the predefined recurrence criterion (detection in ≥3 cancer types). Highlighted nodes and edges denote the subset retained after downstream prioritization and validation procedures, comprising eight miRNAs, 30 target genes, and their corresponding validated associations. The hsa-miR-155-5p–CTLA4–CYT triplet represented the most recurrent positive-track association identified across tumour types. CYT, cytolytic activity score.

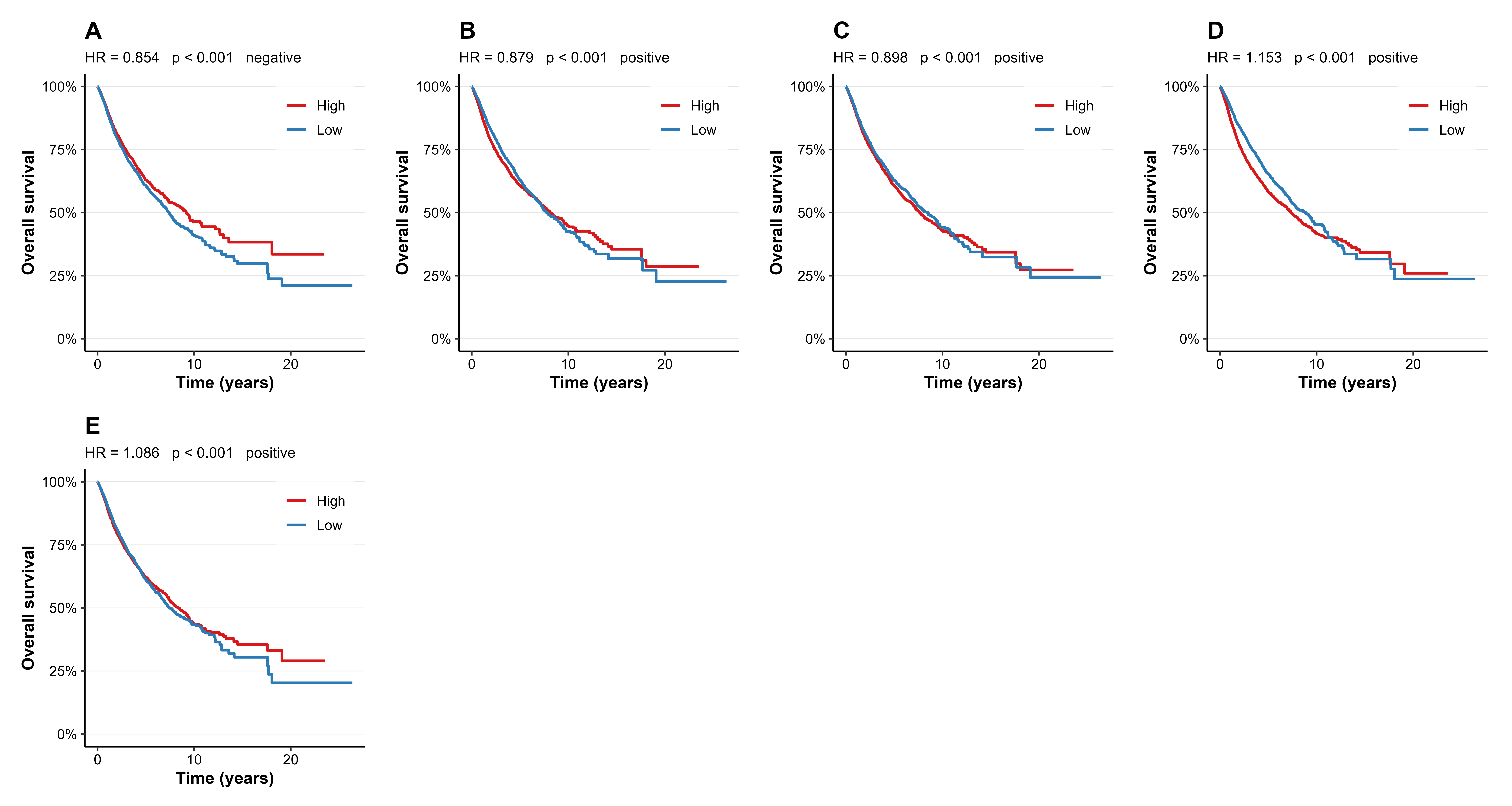

**Supplementary Figure S2. Representative Kaplan–Meier survival analyses of CPE-adjusted candidate miRNAs. (A)** hsa-miR-200b-3p (negative regulatory track), demonstrating a protective association with overall survival. **(B–C)** Protective positive-track miRNAs hsa-miR-146a-5p and hsa-miR-150-5p. **(D–E)** Risk-associated positive-track miRNAs hsa-miR-146b-5p and hsa-miR-130a-3p. Patients were stratified into high- and low-expression groups using the median miRNA expression level. Hazard ratios (HRs) were estimated using cancer-type-adjusted Cox proportional hazards models. Red and blue curves indicate high- and low-expression groups, respectively. Shaded areas represent 95% confidence intervals.
